# The evolutionary assembly of forest communities along environmental gradients: recent diversification or sorting of pre-adapted clades?

**DOI:** 10.1101/2020.12.22.424032

**Authors:** Alexander G. Linan, Jonathan A. Myers, Christine E. Edwards, Amy E. Zanne, Stephen A. Smith, Gabriel Arellano, Leslie Cayola, William Farfan-Ríos, Alfredo F. Fuentes, Karina Garcia-Cabrera, Sebastián Gonzales-Caro, M. Isabel Loza, Manuel J. Macía, Yadvinder Malhi, Beatriz Nieto-Ariza, Norma Salinas Revilla, Miles Silman, J. Sebastián Tello

**Author notes:** These authors contributed equally to this study (Alexander G. Linan, J. Sebastián Tello). corresponding/contact author.

## Abstract

- Biogeographic events occurring in the deep past can contribute to the structure of modern ecological communities. However, little is known about how the emergence of environmental gradients shape the evolution of species that underlie community assembly. In this study, we address how the creation of novel environments lead to community assembly via two non-mutually exclusive processes: 1) the immigration and ecological sorting of pre-adapted clades (ISPC), and 2) recent adaptive diversification (RAD). We study these processes in the context of the elevational gradient created by the uplift of the Central Andes.
- We develop a novel approach and method based on the decomposition of species turnover into within- and among-clade components, where clades correspond to lineages that originated before mountain uplift. Effects of ISPC and RAD can be inferred from how components of turnover change with elevation. We test our approach using data from over 500 Andean forest plots.
- We found that species turnover between communities at different elevations is dominated by the replacement of clades that originated before the uplift of the Central Andes.
- Our results suggest that immigration and sorting of clades pre-adapted to montane habitats is the primary mechanism shaping communities across elevations.

## Introduction

Large-scale biogeographic events—such as the emergence of novel environmental conditions, biotic interchanges, or the evolution of key innovations—can have lasting consequences for biodiversity, community assembly, and species distributions (Ricklefs, 2006; Fussmann *et al.*, 2007; Pelletier *et al.*, 2009; Uribe-Convers & Tank, 2015; Claramunt & Cracraft, 2015). Although theory and empirical evidence suggest that processes occurring in the deep past can contribute to the modern structure of local ecological communities, most research in community ecology during the last few decades has been dominated by a focus on mechanisms at small spatial and temporal scales (Ricklefs, 1987). Studies largely overlook the broader biogeographic context in which communities of co-occurring species are embedded (Chesson, 2000; Adler *et al.*, 2007). Only recently have ecologists begun bridging this gap by developing ecological theory and empirical tests that truly integrate community assembly across eco-evolutionary scales (Emerson & Gillespie, 2008; McGill *et al.*, 2019; Bañares-de-Dios *et al.*, 2020; Segovia *et al.*, 2020). The extent to which community assembly is contingent upon regional context and biogeographic history has broad implications for ecological and evolutionary theory and for understanding how and why communities respond to environmental change (Chase, 2003; Fukami, 2015; Vellend, 2016; McPeek, 2017).

Recent studies provide important insights into how ongoing ecological processes change along environmental gradients (Bricca *et al.*, 2019; Bañares-de-Dios *et al.*, 2020; Neves *et al.*, 2020). However, much less is known about how the emergence of the gradients themselves shape the evolution of species and phenotypes that underlie community assembly. Two non-mutually exclusive processes may explain how communities assemble along gradients following the emergence of novel environmental conditions (Fig. 1). First, the emergence of new environments—e.g., due to climate change, island formation, or mountain orogeny— may create opportunities for immigration and ecological sorting of pre-adapted clades (ISPC hypothesis; Box 1; Fig. 1**a**; Donoghue 2008).

**Figure 1.**
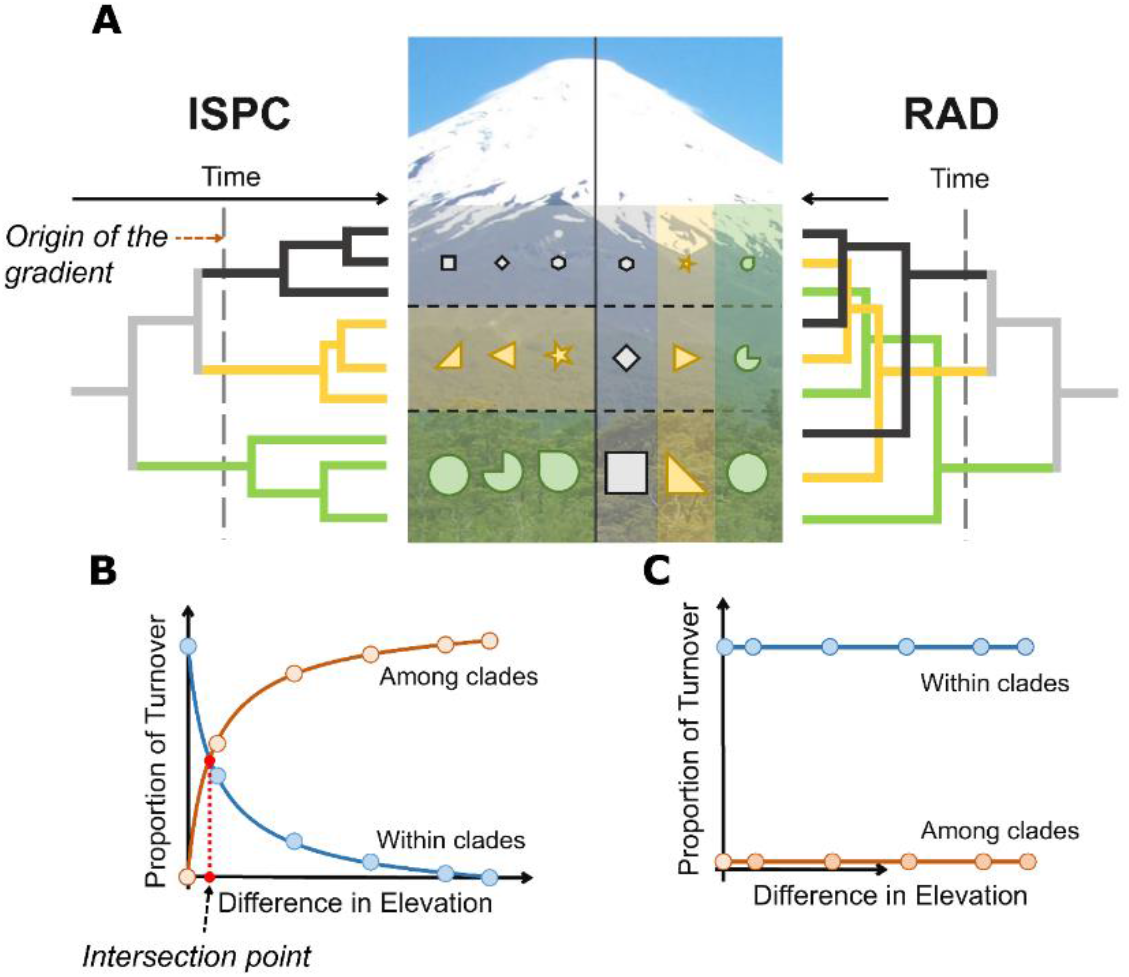
Conceptual models to explain the assembly of regional biotas after the emergence of new environments. (A) shows the distributions of species (symbols), traits (sizes) and clades (colors) along an elevational gradient as expected by the ISPC (left) and RAD (right) hypotheses. The gray broken line marks the emergence of the novel environmental conditions due to mountain uplift. The phylogeny describes the evolutionary relationships among species in the target communities, and the colors indicate different clades of pre-Andean origin (clades that diverged before the uplift of the Central Andes). ISPC and RAD predict contrasting spatial patterns in how species turnover is partitioned into within- and among-clade components. (B) If ISPC is the dominant scenario of community assembly, the among-clade component will increase rapidly as differences in elevation between plots increase, while the within-clade component will decrease. (C) If RAD is the dominant scenario, the within-clade component will dominate the turnover of species as geographic distances increase.

According to this hypothesis, when environmental conditions change within a region and new gradients are created, community assembly across these new habitats is dominated by the immigration of species that are pre-adapted because they occupy similar habitats in a different region. This means that the combination of traits needed to colonize new habitats evolved before the origin of the environmental gradient. Diversification following colonization would not involve adaption to novel environments (i.e. niche conservatism) owing, for example, to competition with species pre-adapted to other environments (Tanentzap *et al.*, 2015; Fukami, 2015). Thus, even though diversification might occur after the origin of the gradient, new species would be restricted mainly to environments to which their ancestors were already pre-adapted. In this way, community assembly across environmental gradients would result in the ecological sorting of species within clades that predate the new environments in the region. This scenario of community assembly is consistent with the idea that “it is easier to move than to evolve” (Donoghue, 2008).

Second, the emergence of new environments may create opportunities for recent adaptive diversification across environments (the RAD hypothesis; Box 1; Fig. 1**a**). According to this hypothesis, when new environmental gradients are created, community assembly across habitats is dominated by adaptation in response to the emerging environmental conditions, resulting in the diversification of clades across the environmental gradient (Stroud & Losos, 2016). Thus, the traits needed to colonize new habitats evolve after the origin of the environmental gradient. In this scenario, niche conservatism is minimal or non-existent, and community assembly results from the diversification of one or more clades that were originally adapted to a subset of environmental conditions, but that diversify to occupy emerging novel environmental space. This scenario for community assembly following the emergence of environmental gradients is consistent with the classic ideas of biome shifts and adaptive radiation driven by ecological opportunity (Schluter, 2000; Losos, 2010; Donoghue & Edwards, 2014).

#### BOX 1 Glossary

##### Pre-Andean clade

A clade that diverged from others before the uplift of the Central Andes. Fig. 1 shows predicted elevational distributions of three pre-Andean clades (colors) based on our hypotheses.

##### Pre-adapted clade

A pre-Andean clade that had, before its immigration to the Central Andes, already evolved adaptations to the novel environmental conditions created by mountain uplift.

##### Turnover

Observed variation in species or functional-trait composition among forest plots or biogeographic regions. For example, in two species assemblages [A, B] & [A, C], turnover is generated by the replacement of species B in the first assemblage with species C in the second.

##### Within-clade turnover

Proportion of total turnover that corresponds to shifts in species composition within a pre-Andean clade. For example, in two assemblages [A, B] & [A, C], within-clade turnover would be high if species B and C belong to the same pre-Andean clade.

##### Among-clade turnover

Proportion of total turnover that corresponds to shifts in species composition among multiple pre-Andean clades. For example, in two assemblages [A, B] & [A, C], among-clade turnover would be high if species B and C belong to different pre-Andean clades.

Here we present and test a novel community-phylogenetic framework and method to disentangle the relative importance of ISPC and RAD in determining the assembly of communities along large-scale environmental gradients. These effects on community assembly can be inferred from unique patterns in the phylogenetic structure of compositional turnover. In particular, signatures of these two processes can be traced when species turnover is decomposed into components that correspond to *within*- and *among-clade turnover*, where clades correspond to independent lineages that originated before the emergence of the gradient. These within- and among-clade turnover components, in turn, reflect the effects of diversification after and before the emergence of the gradient on community composition across environments. Here, we illustrate these patterns using a hypothetical elevational gradient of a mountain depicted in Fig. 1A. The ISPC hypothesis predicts that for communities *at the same elevation*, variation in community composition should be dominated by *within-clade turnover*, reflecting strong niche conservatism of a few clades that are pre-adapted to the environments at that specific elevation (Fig. 1**b**). As communities are *farther apart along the elevational (i.e. environmental) gradient*, variation in community composition should become increasingly dominated by *among-clade turnover*, reflecting the shift in dominance from species in one pre-adapted clade to another. Alternatively, the RAD hypothesis predicts that for communities at *similar or contrasting elevations*, variation in community composition should be dominated by turnover *within clades,* reflecting how multiple clades evolved niche differences in response to new environmental conditions that allow them to have broad elevational (i.e., environmental) distributions (Fig. 1**c**). Although we developed and tested this conceptual framework in the context of mountain uplift, our approach is applicable to study community assembly after the emergence of any type of environmental gradient at any spatial or temporal scale.

In the Neotropics, the geologically recent uplift of the Andean mountains created a striking elevational and environmental gradient that had profound consequences for global climate and biodiversity (Rahbek & Graves, 2001; Antonelli *et al.*, 2009; Ehlers & Poulsen, 2009; Jiménez *et al.*, 2009; Graham, 2009; Hoorn *et al.*, 2010). Indeed, the tropical Andes are considered the most species-rich biodiversity hotspot, containing 15% of all plant species (>45,000 species) in only 1% of the world’s land area (Myers *et al.*, 2000; Rahbek & Graves, 2001; Jiménez *et al.*, 2009). However, our current understanding of the eco-evolutionary forces that shape community assembly across elevations in the hyper-diverse Andean biotas is limited. First, many studies focus on the evolution and distribution of relatively small clades compared to entire communities; these studies have provided evidence for an important role of adaptive diversification (Antonelli *et al.*, 2009; Pérez-Escobar *et al.*, 2017) in some cases and immigration and colonization of pre-adapted clades in others (Hughes & Eastwood, 2006; Jin *et al.*, 2015; Lagomarsino *et al.*, 2016). Such studies demonstrate that both processes have occurred, but provide limited insights into how evolutionary history of individual clades contribute to the assembly of entire ecological communities and regional biotas. Second, studies that focus on the phylogenetic structure of Andean communities are relatively few and often fail to differentiate the effects of diversification before and after the emergence of the gradient (Graham *et al.*, 2009; Parra *et al.*, 2011; Bacon *et al.*, 2018; Montaño-Centellas *et al.*, 2019; Ramírez *et al.*, 2019). To date, no study has sought to disentangle the relative importance of immigration and sorting of pre-adapted clades versus post-Andean uplift adaptive radiation in shaping the enormous variation in plant community composition across elevational gradients.

In this study, we combined tree-species distribution data and phylogenetic information from two large networks of Andean-forest plots to test how RAD and ISPC contribute to the assembly of Andean tree communities. We test these hypotheses in the context of the uplift of the Central Andes, which is associated with the formation of the Altiplano plateau during the last 30 my. (Fig. 1). Moreover, we developed a novel method to decompose measures of species turnover among plots distributed across the elevational gradient into among- and within-pre-Andean clade components (Fig. 1 and Box 1; Legendre & Cáceres 2013). These components measure the relative contributions of ISPC and RAD, respectively. This work provides both a novel framework for examining phylogenetic community turnover and expands our current understanding of how historical processes contribute to community assembly.

## Materials and Methods

### Community composition data across elevations

We utilized data from two large-scale forest plot networks in the Central Andes of Bolivia (the Madidi Project; 30,165 km^2^) and Peru (the Andes Biodiversity and Ecosystem Research Group [ABERG]; 1,765 km^2^). Both datasets contain information on tree community composition spanning the entire elevational range of forests in this region of the Andes from lowland Amazonia to the tree line. Our datasets include information on species composition across 73 1-ha plots (large plots hereafter; 50 in Bolivia and 23 in Peru), as well as 494 0.1-ha plots (small plots hereafter; 458 in Bolivia and 36 in Peru; Fig. 2).

**Figure 2.**
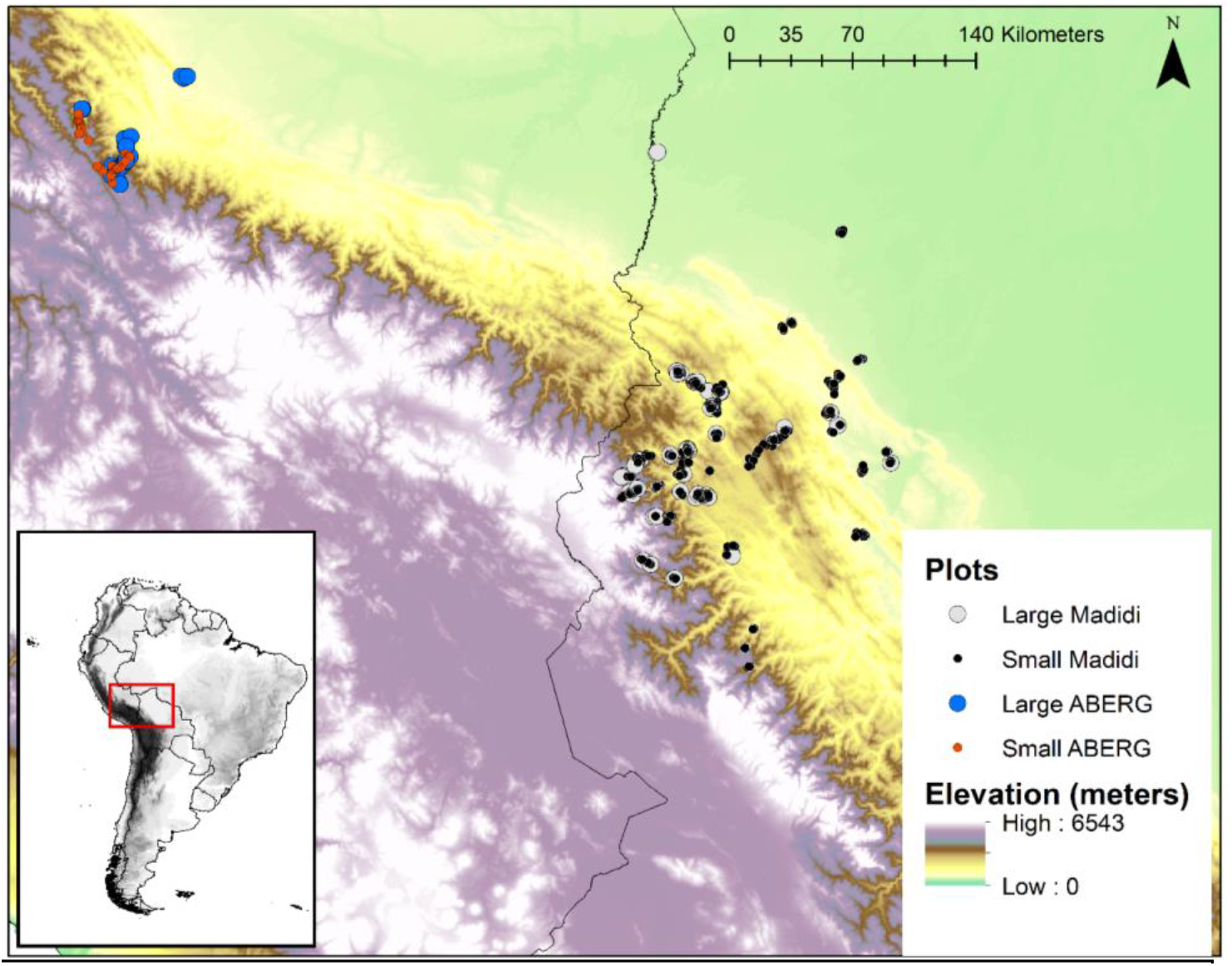
Regional network of forest plots used in this study.

Within these plots, all woody plants with a diameter at breast height (DBH) ≥ 10 cm in large plots and DBH ≥ 2.5 cm in small plots were tagged, measured and identified to species or morpho-species. Large and small plots characterize different plant communities; while large plots consider only adults of large tree species, small plots include younger individuals and also species that 198 do not reach 10 cm DBH, including many shrubs. Thus, we separated out data by plot size and conducted analyses independently. Additionally, we excluded high elevation plots (> 3,800 m) and plots with ≤ 3 species. Most of these plots represent *Polylepis*-dominated forests fragments within a matrix of Paramo grasslands/shrublands. The ecology and composition of these Paramo forests is clearly distinct from the continuous forest cover along the elevational gradient.

Within the Bolivian and Peruvian datasets, we conducted extensive taxonomic work to standardize species and morpho-species names across plots. Morpho-species, however, could not be standardized between the Bolivian and Peruvian data. To test the effect of morphospecies on results, analyses were repeated with and without morphospecies. Both analyses produced nearly identical results (Fig. S1); for simplicity, we present only analyses including morphospecies. Representative specimens at each site were collected and deposited in herbaria, mainly at the Herbario Nacional de La Paz (LPB), the Missouri Botanical Garden (MO) and Universidad Nacional de San Antonio Abad del Cusco (CUZ) in Peru. The final dataset contains 494 small plots and 73 large plots, distributed from 175 m to 4,365 m in elevation. The small plots contained 2,731 species, whereas the large plots contained 1,904 species (Table 1).

**Table 1.**
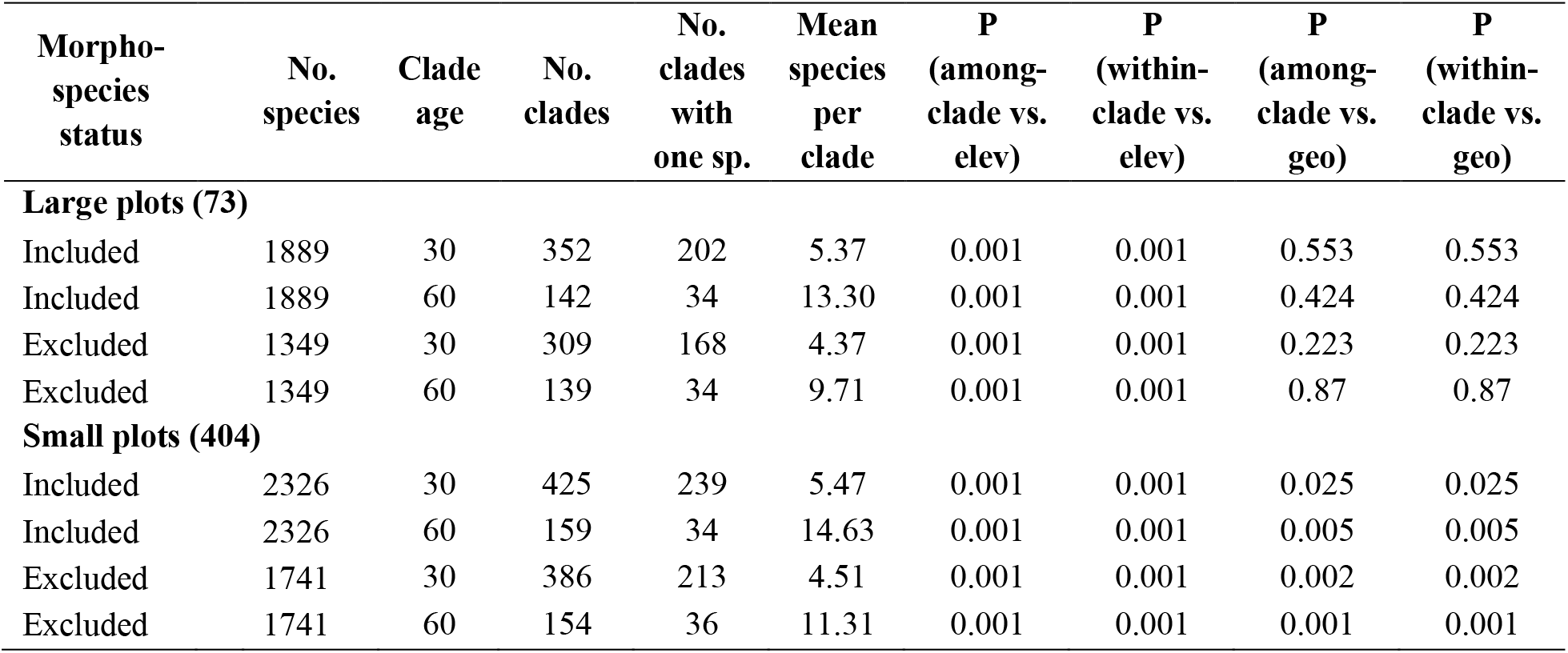
Summary of datasets used for analyses and p-values assessing significance of empirical gradients in among-clade and within-clade turnover across elevational (elev) and geographic (geo) distances.

| Morpho-species status | No. species | Clade age | No. clades | No. clades with one sp. | Mean species per clade | P (among-clade vs. elev) | P (within-clade vs. elev) | P (among-clade vs. geo) | P (within-clade vs. geo) |
| --- | --- | --- | --- | --- | --- | --- | --- | --- | --- |
| **Large plots (73)** |  |  |  |  |  |  |  |  |  |
| Included | 1889 | 30 | 352 | 202 | 5.37 | 0.001 | 0.001 | 0.553 | 0.553 |
| Included | 1889 | 60 | 142 | 34 | 13.30 | 0.001 | 0.001 | 0.424 | 0.424 |
| Excluded | 1349 | 30 | 309 | 168 | 4.37 | 0.001 | 0.001 | 0.223 | 0.223 |
| Excluded | 1349 | 60 | 139 | 34 | 9.71 | 0.001 | 0.001 | 0.87 | 0.87 |
| **Small plots (404)** |  |  |  |  |  |  |  |  |  |
| Included | 2326 | 30 | 425 | 239 | 5.47 | 0.001 | 0.001 | 0.025 | 0.025 |
| Included | 2326 | 60 | 159 | 34 | 14.63 | 0.001 | 0.001 | 0.005 | 0.005 |
| Excluded | 1741 | 30 | 386 | 213 | 4.51 | 0.001 | 0.001 | 0.002 | 0.002 |
| Excluded | 1741 | 60 | 154 | 36 | 11.31 | 0.001 | 0.001 | 0.001 | 0.001 |

### Phylogenetic reconstruction and defining clades of pre-Andean origin

To test our hypotheses, we needed a phylogenetic framework that grouped species into clades that diverged from one-another before the origin of the elevational gradient (i.e. clades that pre-date the uplift of the Central Andes). To do this, we based our analyses on Smith and Brown’s (2018) global mega-phylogeny of seed plants, which combined NCBI sequence data, results from the Open Tree of Life project (Hinchliff *et al.*, 2015), and advances in bioinformatic methods (PyPHLAWD; Smith & Walker 2019) to produce the most comprehensive time-calibrated species-level phylogeny to date. To include species and morphospecies in our dataset that were not in the original phylogeny, we used the R package V.PhyloMaker (Jin & Qian, 2019). Using genus and family level taxonomic information, missing taxa not included in the mega-phylogeny were joined to the halfway point of the family/genus branch (V.PhyloMaker scenario= “S3”). For genera not represented in the mega-phylogeny, we joined species to sister genera in the phylogeny based on support in the literature (when possible) using the ‘bind.relative’ option of V.PhyloMaker. Finally, we pruned from the phylogeny, all species that were absent in our forest plots. The resulting phylogeny included 3,143 species.

The formation of the Andean cordillera has been a complex and heterogeneous process. In the Central Andes, the history of mountain formation is closely tied to the development of the Altiplano plateau, currently located at nearly 3,800 m in elevation. While the traditional view of mountain uplift invokes a slow and gradual process, recent evidence suggests that the uplift of the Altiplano was dominated by spurs of rapid rise with intervening periods of stasis (Garzione *et al.*, 2008, 2017). Although the northern Andes is considered much younger, the best available evidence suggests that most of the uplift in the Central Andes occurred within the last 30 million years. Thus, our analyses used this age as a main reference for the origin of the elevational gradient and to delimit pre-Andean clades.

Pre-Andean clades in the time-calibrated regional phylogeny were defined as those whose stems intersect the 30 my reference. In this way, each pre-Andean clade in our study diverged from others before the uplift of the central Andes, whereas all species within pre-Andean clades resulted from diversification that occurred after mountain uplift had started. We used the function treeSlice in the R package “Phytools” (Revell, 2012) to fragment the regional phylogeny into these clades. Species present in small plots formed 473 pre-Andean clades with an average of 5.77 species per clade, whereas species in the large plots formed 355 clades, averaging 5.36 species per clade (Table 1 and Fig. S2). Finally, we sought see understand how our results varied by defining different ages for pre-Andean clades. Thus, in addition to creating a dataset with pre-Andean clades defined as 30 million years old, we made a second dataset which defined clades as 60 million years old. This represents a much more conservative estimate of the timing of Andean uplift (Hoorn *et al.*, 2010). The results from these alternative analyses were nearly identical, and thus are presented only in the Supplementary Material (Fig. S3).

### Decomposing total turnover into among- and within-clade turnover

To test hypotheses about the relative importance of ISPC and RAD, we developed a method to decompose variation in species turnover between two communities into additive components representing the contribution of turnover among-groups and within-groups (Legendre & Cáceres, 2013). For our analyses, groups are defined by clades of pre-Andean origin, but this decomposition method is broadly applicable to species groupings based on any criteria. Analyses were based on the Sørensen pair-wise dissimilarity index (*S*; Sørensen 1948), which uses presence/absence data:

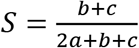

Here, ***a*** represents the number of shared species between two communities, ***b*** is the number of species present only in the first community, and ***c*** is the number of species present only in the second community. Since species are aggregated into clades, species in ***b*** can be further divided into two components: ***b***_WG_ is the fraction of ***b*** that correspond to species in groups present in both communities, while ***b***_AG_ is the fraction of ***b*** corresponding to species in groups present only in the first community. The same process can be done for ***c***, producing the corresponding components ***c***_WG_ and ***c***_AG_. In this way, the additive within-group (*S_WG_*) and among-groups (*S_AG_*) components of Sørensen dissimilarity are defined as:

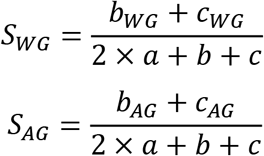

Further details of the decomposition method can be found in the Supplementary Material, where we also show that this approach could be applied to other turnover metrics, such as Bray-Curtis distances. The R code that performs this decomposition is available at https://github.com/Linan552/Madidi-project. When within- and among-clade dissimilarities are transformed into components of total turnover (*S_WG_/S* and *S_AG_/S*, respectively), these values correspond to the contribution of diversification after (*S_WG_*/*S*) and before (*S_AG_*/*S*) the uplift of the central Andes to community species turnover (Fig. 1, S4). Thus, a high among-clade component indicates that turnover is mainly dominated by species that diverged from one another before the uplift of the Central Andes (Fig. 1**a**, left). In contrast, high within-clade component indicates that turnover is dominated by species that diverged from one another after the uplift of the Central Andes (Fig. 1**a**, right).

As described in the introduction, the immigration and sorting of pre-adapted clades (ISPC) and the recent adaptive diversification (RAD) hypotheses make predictions about how these components of turnover will be related to environmental (i.e. elevational) distances. Thus, we plotted the components of turnover for each pair of plots against their elevational distance. The ISPC hypothesis predicts that as communities are farther apart along the elevational gradient, variation in community composition should become increasingly dominated by among-clade turnover (Fig. 1**b**). Alternatively, the RAD hypothesis predicts that variation in community composition should be dominated by turnover within clades regardless of elevational distance (Fig. 1**c**).

### Assessing significance of empirical data using null models and ruling out effects of geographic distance

To test whether observed patterns are different from those expected by chance, we compared the components of turnover in the empirical data with components produced by a null model that eliminated any phylogenetic structure in the distribution of species, but retained other important elements of the data that might shape turnover patters. We ran a “tip-randomization null model” in which species were randomly re-assigned to tips in the phylogeny, such that species were randomly reshuffled among pre-Andean clades. This randomization algorithm maintained the number of species per clade, the diversity gradient across elevations, the average range size in each community, and importantly, the empirical turnover observed between pairs of plots. The only aspect of the data that was randomized was the empirical membership of species in clades of pre-Andean origin. We randomized the data and re-calculated components of turnover for each pair of plots 999 times. From these null expectations, we calculated standardized effect sizes as the empirical value minus the mean of null distribution divided by the standard deviation of the null distribution. These values represent the magnitude of the difference between the empirical components of turnover and the null expectation, where there is no phylogenetic structure in species distributions. As was done for the empirical components of turnover, we related these standardized effect values against the difference in elevation for each pair of plots.

Additionally, we compared the rate of change in turnover components with difference in elevation between the empirical data and null model expectations. To do this, we used slopes from a linear regression between components of turnover and elevational distance. Because these relationships are non-linear, we used a logit transformation on species turnover prior to regression analyses (Cleveland, 1981). These transformations produced a reasonable linearization of the relationships in large and small plot datasets, allowing us to capture the rate of change in a single parameter (see Fig. S5). The empirical slopes were then compared with the distribution of 999 slopes generated by the null model. If empirical slopes were significantly greater or smaller (p<0.05) than null slopes, we concluded that phylogenetic structure exists in the community composition changes along the elevational gradient.

Finally, to control for the effects of geographic distance on the analysis of turnover across elevations, only a subset of pair-wise plot comparisons for each dataset were used. This subset of plot pairs minimized variation in geographic distances, but maximized the elevational range represented in the data (Fig. S6). For the large-plot dataset, we selected plot pairs only between 50 and 90 km apart (8% of the total range in geographic distances), and for the small-plot dataset, we selected plot pairs between 110 and 160 km apart (10% of geographic range). In both datasets, the elevational distances between plots ranged from 175 to 3765 m.

Additionally, we analyzed how the within- and among-clade components of turnover changes as a function of geographic distance. For these analyses, we used pairs of plots spanning the entire geographic range of the study (max distance between plots; 495 km), but were at similar elevations (0 to 200 m of elevational distance, Fig. S6). Like our analyses along elevational distances, we compared patterns of variation in the within- and among-clade turnover against null model expectations. These analyses show how turnover and its components change across space but within the same environmental conditions (results presented in the Supplementary Material).

## Results

Species composition changed dramatically across elevations. Species turnover (Sørensen dissimilarity) among forest plots showed a saturating relationship with elevational distance, increasing rapidly as elevational distance increases and then reaching an asymptote at complete turnover (Fig. 3).

**Figure 3.**
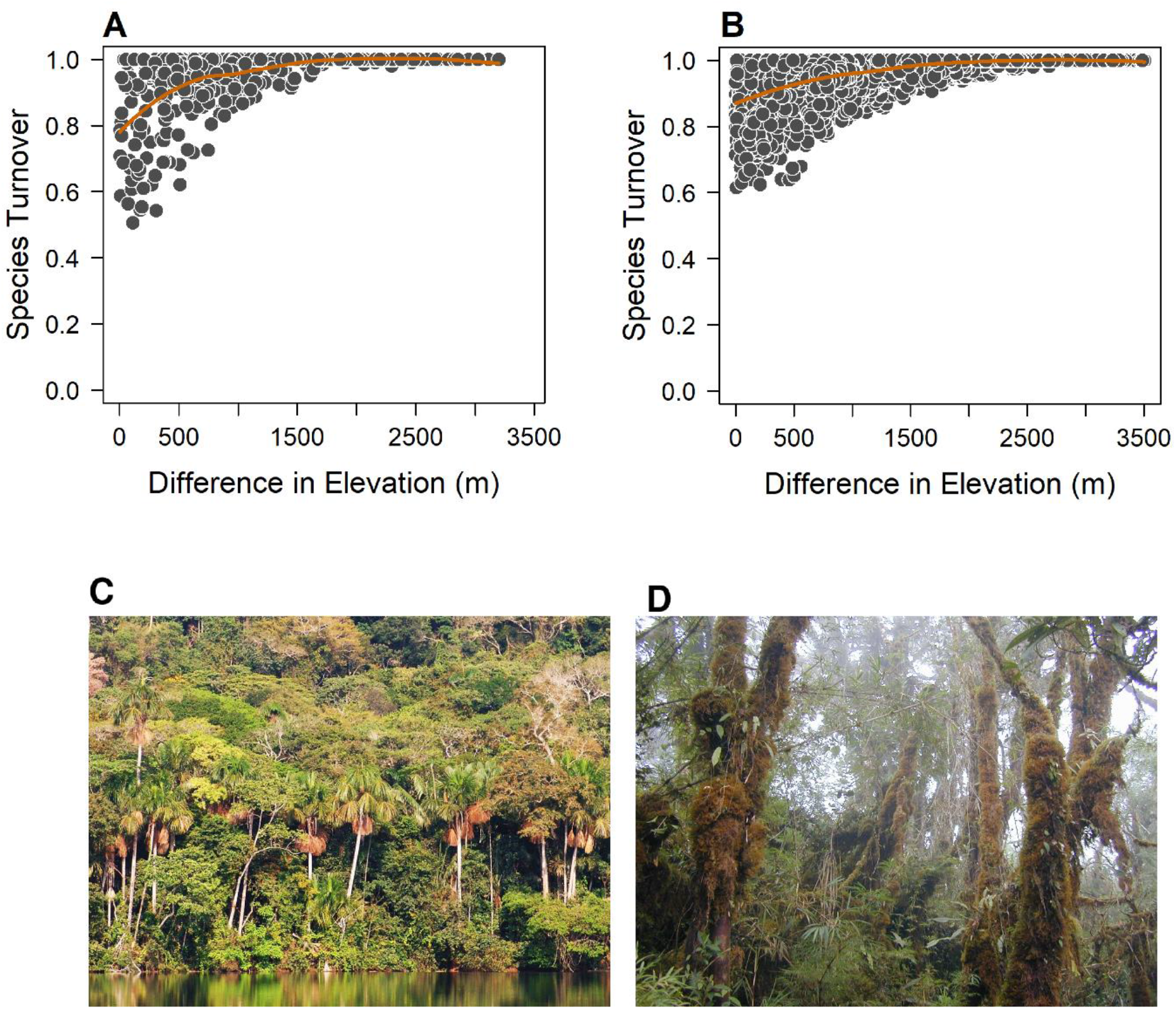
Species turnover across elevations. Sørensen dissimilarity plotted against difference in elevation for each pair of plots in our two datasets. These patterns are presented separately for (A) large 1-ha plots and (B) small 0.1-ha plots. (C; Laguna Chalalan in Bolivia at 400 m elevation) and upper montane cloud forests (D; Trocha Union in Peru at 3,260 m). Pictures by Christopher Davidson, Sharon Christoph and William Farfan-Rios.

Indeed, plots separated by more than 2,000 to 2,500 m of elevation never shared species. We found a similar relationship between species composition and geographic distance (Fig. S7). However, Sørensen dissimilarity did not increase as dramatically with increasing geographic distance and it never reached complete turnover; for example, we found that plots can still share species when they are in similar environments even if plots are 400 km away from one another (one in Peru the other in Bolivia; Fig. S7).

We also found strong elevational gradients in the within- and among-clade components of species turnover (Fig. 4**a,d**). For forest plots occurring at the same elevation (zero meters in elevational difference), among- and within-clade components were equal in magnitude (Fig. 4**a,d**). This result indicates that communities in the same environment shared species in the same pre-Andean clades, but also that multiple different clades contributed to community composition among these plots. As elevational difference increased, among-clade turnover rose rapidly, while within-clade turnover decreased (Fig. 4**a,d**). Although the increase in the among-clade component was monotonic in the small-plots dataset, it saturated at around 2,000 m of elevational difference for the large-plots dataset. In both datasets, however, when plots were separated by more than 1,000 to 1,500 m in elevation, pairs of communities were found where 100% of the turnover corresponded to the among-clade component. This result means that some pairs of plots at opposite ends of the elevational gradient shared neither species nor clades 30 my old, which originated before the uplift of the Central Andes. These results support the idea that the immigration and ecological sorting of pre-Andean clades had a major effect in shaping community composition across elevations - the ISPC hypothesis.

**Figure 4.**
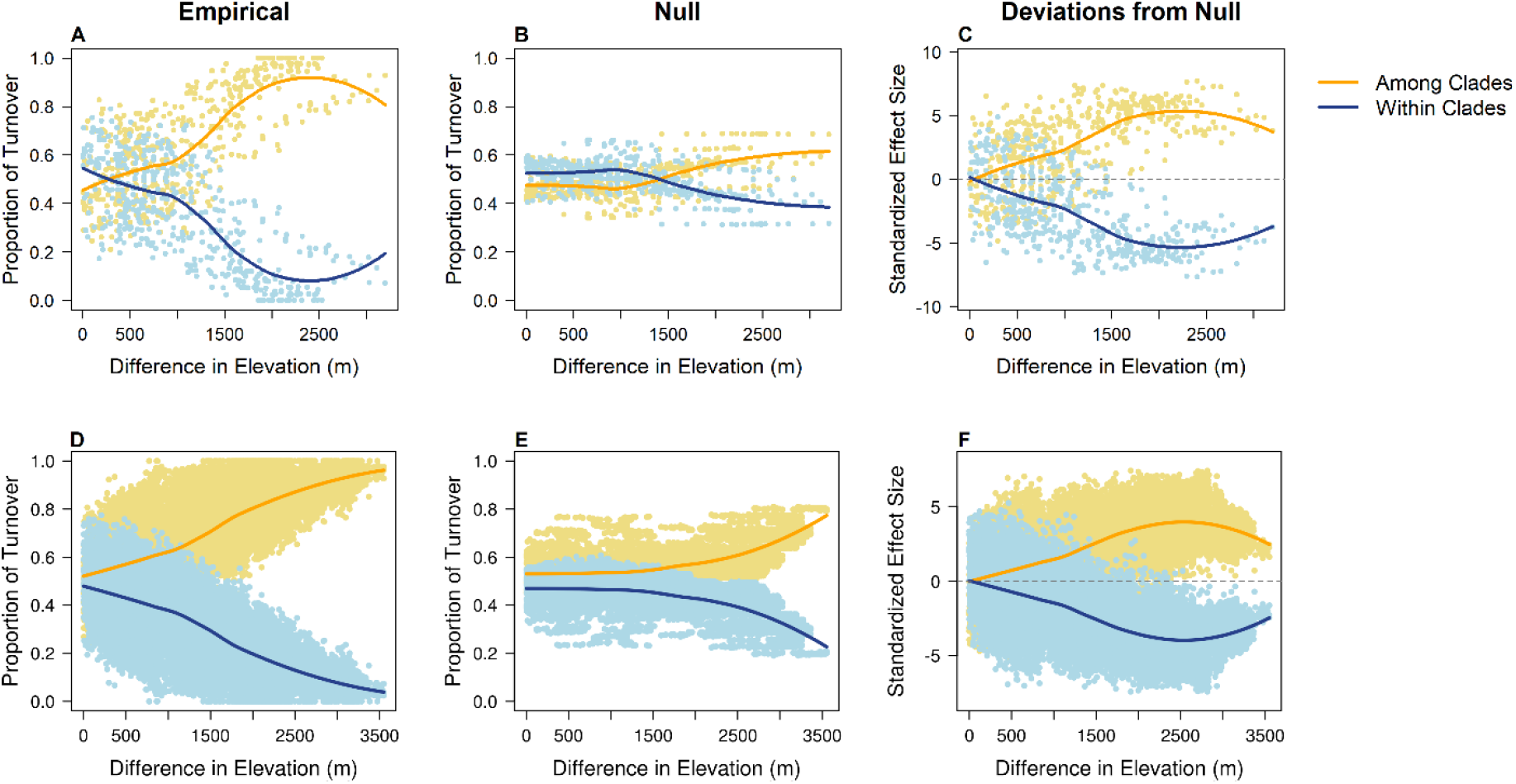
Decomposition of species turnover across elevational gradients into among-clade and within clade components – 30 MY clades from small and large plots. Sorensen dissimilarities between each pair of plots were decomposed into within-clades (blue lines) and among-clades (yellow lines) components. We then plotted these components of turnover against difference in elevation in large plots (first row) and small plots (second row). Finally, we compared spatial patterns in variation of these components with a tip-randomization null model that removes any phylogenetic structure in the distribution of species across elevation. (A. & D.) empirical patterns; (B. & E.) patterns for the mean of the expectations in the null model; (C & F) patterns for standardized effect sizes showing the deviation of the empirical values from null expectations.

The predictions of the ISPC hypotheses were also supported using standardized effect sizes – as measured using our null model (Fig. 4 **c,f**). Indeed, standardized effect sizes for within- and among-clade components were both close to zero for plots at the same elevation. As elevational differences increased, standardized effect sizes increased for among-clade turnover and decreased for within-clade turnover (Fig. 4 **c,f**). Moreover, when plots were separated by >1,500 m in elevation, the empirical values differed by more than two standard deviations from null expectations (i.e. standardized effect sizes greater than 2; Fig. **c,d**). The comparison of the empirical and null regression slopes also showed that the change in the empirical components was much more pronounced than the change expected by the null model (Fig. 5).

**Figure 5.**
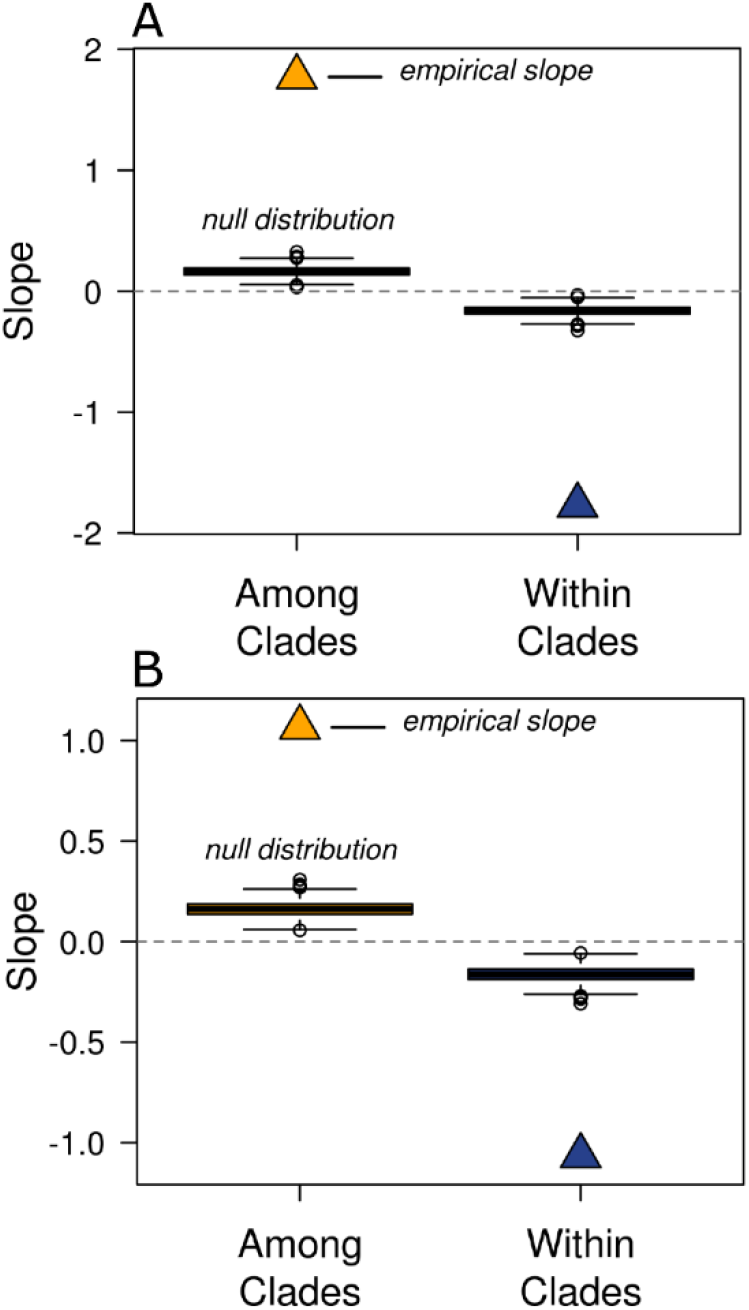
Comparison of linear slopes. Empirical slopes of logit transformed proportional turnover across elevational distance (triangles) compared to the distribution of 999 slopes produced by the null model (boxplots) in (A) large plots and (B) small plots.

Finally, we found that geographic distance did not have the same effect on components of turnover as elevational difference. Among large plots, among- and within-clade turnover remained constant and of similar magnitude with increasing geographic distance (Fig. 4 vs. S8). For small plots, on the other hand, the magnitude of the within-clade turnover component increases with geographic distance. This pattern remained when using standardized effect sizes (Fig. 4 vs. S8).

## Discussion

### Community assembly across contrasting elevations is dominated by the immigration and ecological sorting of clades that pre-date mountain uplift

Our results showed clearly that changes in species composition across elevations were driven primarily by a replacement of clades of pre-Andean origin. These results were robust to analyses using different age estimates of pre-Andean clades (30 vs. 60 mya), inclusion or exclusion of morpho-species, or delimitations of forest communities (trees ≥ 10 cm DBH in large plots vs. trees ≥ 2.5 cm DBH in small plots). While adaptive diversification is likely to have occurred in our study system, our results suggest that this process has had a reduced influence on patterns of community assembly. In contrast, we found strong evidence for a high relative importance of the ISPC hypothesis. The new environments created by the uplift of the Central Andes during the last 30 my were colonized primarily by clades of species that were pre-adapted to the emerging environmental conditions. Diversification within these clades resulted in new tree species that had elevational distributions similar to those occupied by the immigrating species. In this way, the ecological sorting of pre-Adapted clades according to their pre-adaptations is the eco-evolutionary process that dominates the regional assembly of tree communities across the elevational gradient.

Our study focuses on the structure of species assemblages, and how biogeographic processes shape patterns of diversity. The assembly of communities, however, integrates the evolutionary history of multiple independent clades of species. Several previous studies that have taken this approach, focusing on the evolution of clades after the Andean uplift. This research shows that groups of animals and plants across the Andes have diversified in ways that are consistent with our results (Bell & Donoghue, 2005; Hughes & Eastwood, 2006; Chaves *et al.*, 2011; Nürk *et al.*, 2013). One of the best studied biogeographic histories in the Andes is that of the plants in the genus *Lupinus*, which colonized the Andes from temperate North America (Hughes & Eastwood, 2006), and were likely pre-adapted to the cold conditions of alpine environments (Nevado *et al.*, 2016). This clade experienced an explosive diversification in the Andes, but most of the resulting species occupy only high-elevation habitats. Their diversification was likely fueled by the interaction between insularity of high-mountain habitats and climatic fluctuations during Quaternary (Nevado *et al.*, 2018). Adaptive diversification also played an important role in the radiation of the Andean lupins (Nevado *et al.*, 2016). Indeed, species in the clade show a huge diversity of phenotypes, life forms and micro-habitat use (Hughes & Eastwood, 2006). Their adaptive diversification, however, did not involve large numbers of species colonizing the new environments at different elevations created by mountain uplift. Several clades of plants distributed at the highest elevations in the Andes seem to show similar patterns of diversification (Madriñán *et al.*, 2013). A general pattern of conservatism in elevational distribution was also documented for several clades of trees by Griffiths et al. (2020). Clades with a biogeographic history similar those of the Andean lupins would contribute little to changes in species composition along the elevational gradient. Instead, this pattern of diversification, when experienced by numerous clades pre-adapted to different elevations can lead to the observed patterns of clade turnover found in our study. Indeed, turnover among communities across the elevational gradient have an evolutionary origin that is rooted deep in the past, and that mostly pre-dates the emergence of the environmental gradient.

Studies of particular clades, like those highlighted above, are insightful and have helped us advance our understanding of the patterns and mechanisms of diversification. However, this approach does not address directly the eco-evolutionary forces behind the assembly of diverse communities, which is the focus of our analyses. To the best of our knowledge, our study is the first effort to explicitly test the role that diversification before and after the origin of the environmental gradient (i.e., the uplift of the Central Andes) had on community structure across elevations. While previous studies have not tested the role of mountain uplift directly, our results are supported by previous research of Andean communities, which have suggested an important role for niche conservatism in community assembly across elevations (Graham *et al.*, 2009; Hardy *et al.*, 2012; Jin *et al.*, 2015; Ramírez *et al.*, 2019; Worthy *et al.*, 2019; Bañares-de-Dios *et al.*, 2020). A recent important study in this respect is that by Segovia *et al.* (2020), who demonstrated a clear link in the phylogenetic composition of Andean tree communities to temperate regions of North and South America. In particular, they highlight the role that freezing conditions at high elevations play in creating environments that are invaded by temperate clades. Similarly, niche conservatism has been implied in the eco-evolutionary assembly of seasonally dry forest communities, which occur in rain-shadowed valleys along the Andes. Our study, however, goes further than simply demonstrating niche conservatism or phylogenetic clustering of communities. Instead, we provide evidence that the assembly of communities across elevations is primarily driven by the immigration and sorting of clades that evolved appropriate adaptations even before the emergence of the environmental gradient (Hardy *et al.*, 2012; Chi *et al.*, 2014; Kubota *et al.*, 2018).

### Mountain uplift might create opportunities for adaptive radiation, but this process has a limited effect on community assembly along elevational gradients

Adaptive radiations have played a critical role in the formation of biodiversity, giving rise to an often-stunning array of morphological and species diversity (Gillespie *et al.*, 2020). Previous studies have suggested that ecological opportunity is an important determinant, maybe a prerequisite, of adaptive radiations (Stroud & Losos, 2016; Gillespie *et al.*, 2020), allowing species to diversify rapidly to fill available niche space. The uplift of the Central Andes created environments that were previously unavailable in the region, likely opening up new unoccupied niche space for species. Moreover, as we discussed earlier, numerous rapid radiations have been documented in the Andes (Madriñán *et al.*, 2013); some of them, like that of *Lupinus* or *Espeletia* (Hughes & Eastwood, 2006; Pouchon *et al.*, 2018) are as dramatic as those in clades that epitomize adaptive radiation (e.g. stickleback fish or African Great Lake cichlids; Gillespie *et al.* 2020). If ecological opportunity existed and rapid diversification in the mountains is well documented, then why did we not find a strong signal for recent adaptive radiation in the assembly of communities across elevations?

There are several reasons that could explain our lack of evidence for recent adaptive radiations across the elevational gradient. First, recent and rapid radiations in the Andes may not involve adaptive diversification. Instead, high rates of species accumulation could be fueled solely by allopatric speciation resulting from repeated cycles of habitat isolation and reconnection driven by climatic oscillations (Nevado *et al.*, 2018; Flantua *et al.*, 2019). This process would produce a large number of species that replace one another across geography but within the same environment (Hughes & Eastwood, 2006; Chaves *et al.*, 2011). Furthermore, nearly all cases of recent montane radiations occur in clades occupying the highest elevations (Bell & Donoghue, 2005; Hughes & Eastwood, 2006; Nürk *et al.*, 2013; Hughes & Atchison, 2015), and more studies are needed to know if these patterns of diversification also occur at mid or low elevations (but see Lagomarsino *et al.* 2016). Second, adaptive radiations may have occurred along environmental dimensions other than those of the elevational gradient. Indeed, some of the classic examples of Andean diversification involve fast evolution of phenotypes, even if the elevational distribution of the clade is highly conserved (Hughes & Eastwood, 2006; Nürk *et al.*, 2018; Pouchon *et al.*, 2018). Finally, some clades may have adaptively radiated across the elevational gradient, but these clades are rare and contribute little to overall assembly patterns. Indeed, biogeographic studies have documented significant shifts in elevational distribution during the evolutionary history of several groups of plants and animals (Elias *et al.*, 2009; Bacon *et al.*, 2018). Some of these shift in elevational distribution might be accompanied by shifts in life form (as exemplified by *Espeletia*; Pouchon *et al.* 2018; but see Zanne *et al.* 2013) which do not contribute to the assembly of tree communities that are the focus of our study. The relative frequency of adaptive diversification across elevations versus niche conservatism has not been evaluated; however, Griffith *et al.* (2020) found that in Peru, most clades of trees have narrow elevational distributions, and only a few show wider elevational distributions than expected by chance. The overall lack of clades that appear to have adaptively radiated across elevations could be due to evolutionary priority effect, whereby different elevational niches may have been preempted by immigrating pre-adapted clades (Fukami, 2015). While the role of adaptive radiation in community assembly deserves further study, our results suggest that Andean community assembly is mainly the result of different pre-adapted clades that originated before Andean uplift, which colonized available niches before other clades could adaptively radiate to occupy a broad elevational gradient (Tanentzap *et al.*, 2015).

### Conclusions: future directions and implications for conservation

In this study, we developed a novel conceptual framework (Fig. 1), as well as new methods of decomposing species turnover (Fig. 1, S4), to investigate the biogeographic origins of community assembly along environmental gradients. We use this approach to study how the uplift of the Central Andes led to the variation in community composition along iconic elevational gradients. Our approach, however, can be applied to any system in which the timing of the emergence of an environmental gradient is known and time-calibrated phylogenies can be generated. We envision future studies using this method to understand the eco-evolutionary assembly in systems beyond Andean forests such as along precipitation gradients (Parolari *et al.*, 2020), across contrasting soil conditions (Capurucho *et al.*, 2020), or even under different disturbance regimes (Cavender-Bares & Reich, 2012). Methods such as these can be used to test hypotheses about specific process of community assembly, going beyond documenting niche conservatism or phylogenetic aggregation. Our approach will facilitate deeper insights into how the emergence of environmental gradients shape modern natural ecosystems.

Our analyses demonstrate that species turnover across elevations in the Central Andes is driven primarily by the turnover of clades that are at least 30 my old. These results suggest a strong role for immigration and ecological sorting of pre-adapted clades to the novel environments across elevations created by the uplift of the Central Andes. Adaptive diversification following the emergence of the elevational gradient is likely restricted to a few clades or to narrow elevational bands, having little impact on the assembly of communities along such a large environmental gradient. Our results add to a growing body of evidence suggesting that present day communities are strongly influenced by the ability of lineages to track environmental conditions through space and geological time (Emerson & Gillespie, 2008; Donoghue, 2008; Carvajal-Endara *et al.*, 2017; Griffiths *et al.*, 2020; Segovia *et al.*, 2020).

This finding has important implications for the long-term persistence of communities facing the effects of human-mediated global change. Increases in atmospheric temperatures are predicted to cause elevational shifts in environmental conditions, such that climates that currently occur at specific elevations will occur at higher elevations in the future (Harsch *et al.*, 2009; Ruiz-Labourdette *et al.*, 2012; Freeman *et al.*, 2018; O’Sullivan *et al.*, 2020). Our work on historical patterns of community assembly suggests that ecosystems are more likely to track shifting habitats rather than adapt to novel conditions (Sheldon *et al.*, 2011; Ruiz-Labourdette *et al.*, 2012; Freeman *et al.*, 2018; Feeley *et al.*, 2020). Communities and species at the highest elevations might be specially threatened by climate change since their environments will disappear at the top of mountains and new pre-adapted competitors will move in from lower elevations (Colwell *et al.*, 2008). Thus, communities occupying the highest-elevation sites in the Andes should be prioritized for monitoring and conservation efforts. Because their habitat may not persist over the long term, ex situ conservation (either through conservation seed banking or living collections) of the species endemic to the highest elevations should be a specific priority.

## Acknowledgements

We thank the Dirección General de Biodiversidad, the Bolivian Park Service (SERNAP), the Madidi National Park and local communities for permits, access, and collaboration in Bolivia, where fieldwork was supported by the National Science Foundation (DEB 0101775, DEB 0743457, DEB 1836353). Additional financial support to The Madidi Project has been provided by the Missouri Botanical Garden, the National Geographic Society (NGS 7754-04 and NGS 8047-06), the Comunidad de Madrid (Spain), Consejo Superior de Investigaciones Cientiíficas (Spain), Centro de Estudios de Ameérica Latina (Banco Santander and Universidad Autoónoma de Madrid, Spain), and the Taylor and Davidson families. Fieldwork in the ABERG transect was supported by NSF, the Gordon and Betty Moore Foundation and the UK Natural Environment Research Council. We thank all the researchers, students and local guides that were involved in the collection of the data, particularly Carla Maldonado, Maritza Cornejo, Alejandro Araujo, Javier Quisbert, Narel Paniagua and Peter Jørgenson. Finally, we thank Iván Jiménez for helpful discussions, ideas and comments.

## Author contribution

JST, JAM, AEZ and CEE developed and designed the study. JST, CL, AFF, MIL, GA and MJM collected the Madidi Project dataset; MS, WFR, KGC, NSR, and YM collected the ABERG dataset. SAS produced the phylogenetic data. AGL and JST performed data analyses. AGL and JST wrote the manuscript, and all authors contributed significantly to revisions.

## Data availability

The Madidi Project’s dataset used in our analyses correspond to version 4.1, which is deposited in Zenodo (https://doi.org/10.5281/zenodo.4276558). Additionally, raw data of the Madidi Project are stored and managed in Tropicos (https://tropicos.org/home), the botanical database of the Missouri Botanical Garden. These data can be viewed and accessed via the Madidi Project’s module at http://legacy.tropicos.org/Project/MDI. The Andes Biodiversity and Ecosystem Research Group (ABERG) is a team of 38 researchers from 12 universities dedicated to understanding biodiversity distribution and ecosystem function in the Peruvian Andes. ABERG is committed to data exchange within the scientific community and promoting collaboration among other tropical ecosystem scientists. For more information and to request data contact Miles Silman or Yadvinder Malhi (http://www.andesconservation.org/). The R code created for analyses is available at https://github.com/Linan552/Madidi-project.

## Supporting Information legends

**Supplementary methods: Additive decomposition of species turnover metrics.**

**Figures S1: decomposed turnover across elevation excluding morphospecies.**

**Figures S2: number of species per clade across datasets.**

**Figure S3: decomposition of species turnover across elevational distance for 60my clades.**

**Figures S4: decomposed turnover across elevation excluding morphospecies.**

**Figure S5: logit transformed decomposition of turnover.**

**Figure S6: relationship between difference in elevation and geographic distance in our datasets.**

**Figure S7: Species turnover across geographic distance.**

**Figure S8: Decomposition of species turnover across geographic distance for 30my clades.**

