## Supplemental Information for "The evolutionary assembly of forest communities along environmental gradients: recent diversification or sorting of pre-adapted clades?"

**Supplemental Information: Andean uplift and drivers of community assembly: recent diversification or sorting of pre-adapted clades?**

***Table of Contents***

| **Supplementary methods: Additive decomposition of species turnover metrics** | Page 1 |
| --- | --- |
| **Figures S1: decomposed turnover across elevation excluding morphospecies** | Page 5 |
| **Figures S2: number of species per clade across datasets** | Page 6 |
| **Figure S3: Decomposition of species turnover across elevational distance 60my clades** | Page 7 |
| **Figures S4: decomposed turnover across elevation excluding morphospecies** | Page 8 |
| **Figure S5: Logit transformed decomposition of turnover** | Page 9 |
| **Figure S6: Relationship between difference in elevation and geographic distance in our datasets** | Page 10 |
| **Figure S7: Species turnover across geographic distance** | Page 11 |
| **Figure S8: Decomposition of species turnover across geographic distance 30my clades** | Page 12 |

***Supplementary Methods: Additive decomposition of species turnover metrics (Sørensen and Bray-Curtis dissimilarities) into within- and among-group components***

Turnover (i.e. beta-diversity) is a complex measure of how community structure changes through space and time (Anderson *et al.* 2011). Turnover results from co-varying patterns in the distribution and abundance of multiple species. As such, a multitude of metrics exists to estimate turnover, each of which can reflect different aspects of variation community composition (Tuomisto 2010b, a). Several turnover metrics can also be partitioned into components that might represent different sub-structures in community change (e.g. richness vs. replacement; Baselga & Leprieur 2015), or that correspond to the contribution of individual species, individual sites or groups of sites (Legendre & Cáceres 2013). Partitioning turnover can provide important insights into the forces shaping community composition by allowing us to study how different facets of turnover respond to observational or experimental factors. In this study, we provide a novel mathematical decomposition of pairwise-turnover metrics into additive components that correspond to within- and among-groups contributions. This decomposition allows us to study how diversification before and after a specific event (in our case the uplift of the Central Andes) shape community structure across an environmental gradient.

Unlike the approach developed by Legendre and Caseres (2013); our decomposition is based on groups of species, not sites, and occurs at the level of individual pair-wise dissimilarities, rather at the level of the full species-by-sites matrix. We develop this decomposition for Sørensen and Bray-Curtis dissimilarity metrics, but it could potentially be extended to other metrics as well. Moreover, while the species groups in our study are defined by clades of pre-Andean origin, this method can be used with any criteria to aggregate species into groups. To our knowledge, this is the first time that such a decomposition has been used. We also provide R code with a function to carry out the decomposition outlined here.

The traditional formula for Sørensen dissimilarity (*S*) is

$S=1-\frac{2a}{2a+b+c}$(Legendre & Legendre 2012)

which can be re-written as

$$S=\frac{b+c}{2a+b+c}$$

Here, ***a*** represents the number of shared species between two communities, ***b*** is the number of species present in the first community, but absent in the second community, and ***c*** is the number of species absent in the first community, but presents in the second community. If species are aggregated into groups (by clades, for example, Figure S1A), then ***b*** can be divided into components so that ***b*** = ***b***_WG_ + ***b***_AG_. Here ***b***_WG_ is the fraction of ***b*** that correspond to species in shared groups (i.e. groups that have at least one representative in both communities), while ***b***_AG_ is the fraction of ***b*** that correspond to species in groups unique or endemic to the first community (i.e. groups that have representatives only in the first community). In this way, ***b***_WG_ represents species turnover within groups, and ***b***_AG_ represents species turnover among groups. The same decomposition can be done focusing on the second community so that ***c*** = ***c***_WG_ + ***c***_AG_. In Figure S1, we illustrate an example of how b and c values as well as Sørensen dissimilarities can be partitioned into within- and among-group components. In this way, the formula for Sørensen dissimilarity can be rewritten as:

$$S=\frac{\left( b_{WG}+b_{AG} \right)+\left( c_{WG}+c_{AG} \right)}{2\times a+b+c}$$

$$S=\frac{b_{WG}+c_{WG}}{2\times a+b+c}+\frac{b_{AG}+c_{AG}}{2\times a+b+c}$$

Then, if we define

$$S_{WG}=\frac{b_{WG}+c_{WG}}{2\times a+b+c}$$

$$S_{AG}=\frac{b_{AG}+c_{AG}}{2\times a+b+c}$$

*S* can be partitioned in the two additive components: species turnover within clades (*S_WG_*) and species turnover among clades (*S_AG_*):

$$S=S_{WG}+S_{AG}$$

Although it was not part of our main analyses, we also developed an additive partitioning of the abundance-based Bray-Curtis dissimilarity (also known as the percent difference or the Steinhaus dissimilarity; (Legendre & Legendre 2012). The formula for Bray-Curtis dissimilarities is:

$BC=1-\frac{2\times W}{A+B}$ (Legendre & Legendre 2012)

Here, ***W*** is the sum of the minimum abundances of each species across the two communities; ***A*** is the total sum of abundances in the first community and ***B*** is the total sum of abundances in the second community (Legendre & Legendre 2012). The formula can be re-written as:

$$BC=\frac{A+B-2\times W}{A+B}$$

Like for Sørensen dissimilarities, if species are aggregated into groups (Figure S1A), then the *A*, *B* and *W* values can be divided into fractions. For example, the value *A* (the abundance of species in the first community) can be calculated for three sets of species so that *A = A_WS_* + *A_WG_* + *A_AG_*. Because *BC* uses abundance data, ***A_WS_*** is calculated for species that are present in both communities, but vary in abundance. Thus, *A_WS_* represents within-species turnover. *A_WG_* and *A_AG_* are both calculated for species that are present in community 1, but absent in community 2. *A_WG_*, however, uses only species in shared groups (groups that have species in both communities), while *A_AG_* uses species in groups that only occur in community 1. Thus, *A_WG_* corresponds to species turnover within groups, and *A_AG_* to species turnover among groups. Similar calculations can be done for *B* and *W*, so the formula for *BC* can be re-written as:

$$BC=\frac{\left( A_{WS}+A_{WG}+A_{AG} \right)+\left( B_{WS}+B_{WG}+B_{AG} \right)-2\times\left( W_{WS}+W_{WG}+W_{AG} \right)}{A+B}$$

$$BC=\frac{A_{WS}+B_{WS}+2\times W_{WS}}{A+B}+\frac{A_{WG}+B_{WG}+2\times W_{WG}}{A+B}+\frac{A_{AG}+B_{AG}+2\times W_{AG}}{A+B}$$

The *W* fractions represent the sums of minimums across sites. Thus, *W_WG_* and *W_AG_* become zero because they are based on species present only in one site (i.e. minimum abundances of 0 for all species).

$$BC=\frac{A_{WS}+B_{WS}+2\times W_{WS}}{A+B}+\frac{A_{WG}+B_{WG}}{A+B}+\frac{A_{AG}+B_{AG}}{A+B}$$

If we define:

$${BC}_{WS}=\frac{A_{WS}+B_{WS}+2\times W_{WS}}{A+B}$$

$${BC}_{WG}=\frac{A_{WG}+B_{WG}}{A+B}$$

$${BC}_{AG}=\frac{A_{AG}+B_{AG}}{A+B}$$

*BC* can be partitioned in the three additive components: turnover within species (*BC_WS_*), species turnover within clades (*BC_WG_*) and species turnover among clades (*BC_AG_*):

$$BC={BC}_{WS}+{BC}_{WG}+{BC}_{AG}$$

***Supplementary Material References***

Anderson, M.J., Crist, T.O., Chase, J.M., Vellend, M., Inouye, B.D., Freestone, A.L., *et al.* (2011). Navigating the multiple meanings of β diversity: a roadmap for the practicing ecologist. *Ecol. Lett.*, 14, 19–28.

Baselga, A. & Leprieur, F. (2015). Comparing methods to separate components of beta diversity. *Methods Ecol. Evol.*, 6, 1069–1079.

Legendre, P. & Cáceres, M. De. (2013). Beta diversity as the variance of community data: dissimilarity coefficients and partitioning. *Ecol. Lett.*, 16, 951–963.

Legendre, P. & Legendre, L. (2012). *Numerical ecology*. Elsevier.

Tuomisto, H. (2010a). A diversity of beta diversities: straightening up a concept gone awry. Part 1. Defining beta diversity as a function of alpha and gamma diversity. *Ecography (Cop.).*, 33, 2–22.

Tuomisto, H. (2010b). A diversity of beta diversities: straightening up a concept gone awry. Part 2. Quantifying beta diversity and related phenomena. *Ecography (Cop.).*, 33, 23–45.

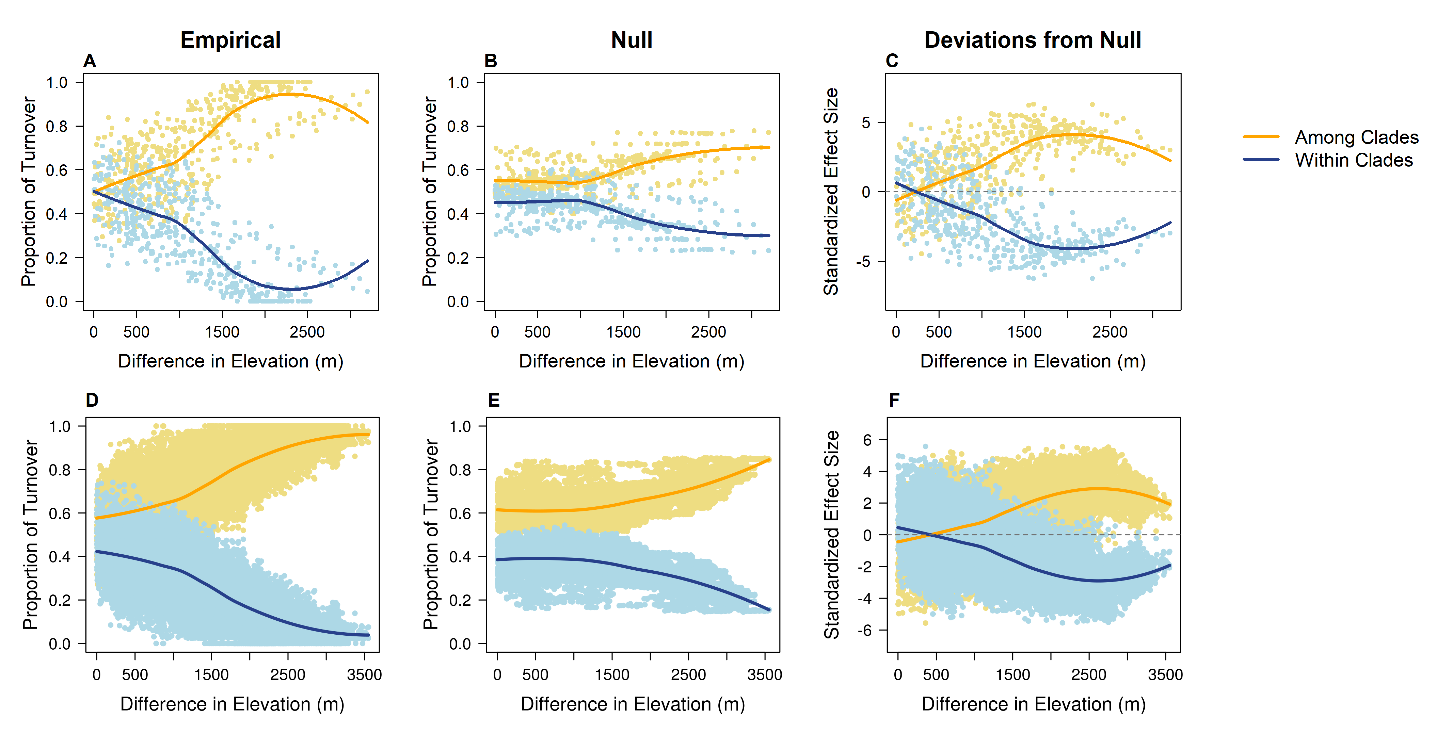

**Figure S1.** **Decomposition of species turnover across elevational distance into among-clade and within-clade components – 30 MY clades from small and large plots excluding morphospecies.** Sørensen dissimilarities between each pair of plots were decomposed into within-clades (blue lines) and among-clades (yellow lines) proportions. We then plotted these components of turnover against elevational distance in large plots (first row) and small plots (second row). Finally, we compared spatial patterns in variation of these components with a tip-randomization null model that removes any phylogenetic structure in the distribution of species across elevational distance. **(A & E**) empirical patterns; **(B & E)** patterns for the mean of the expectations in the null model; **(C & F)** patterns for standardized effect sizes showing the deviation of the empirical values from null expectations.

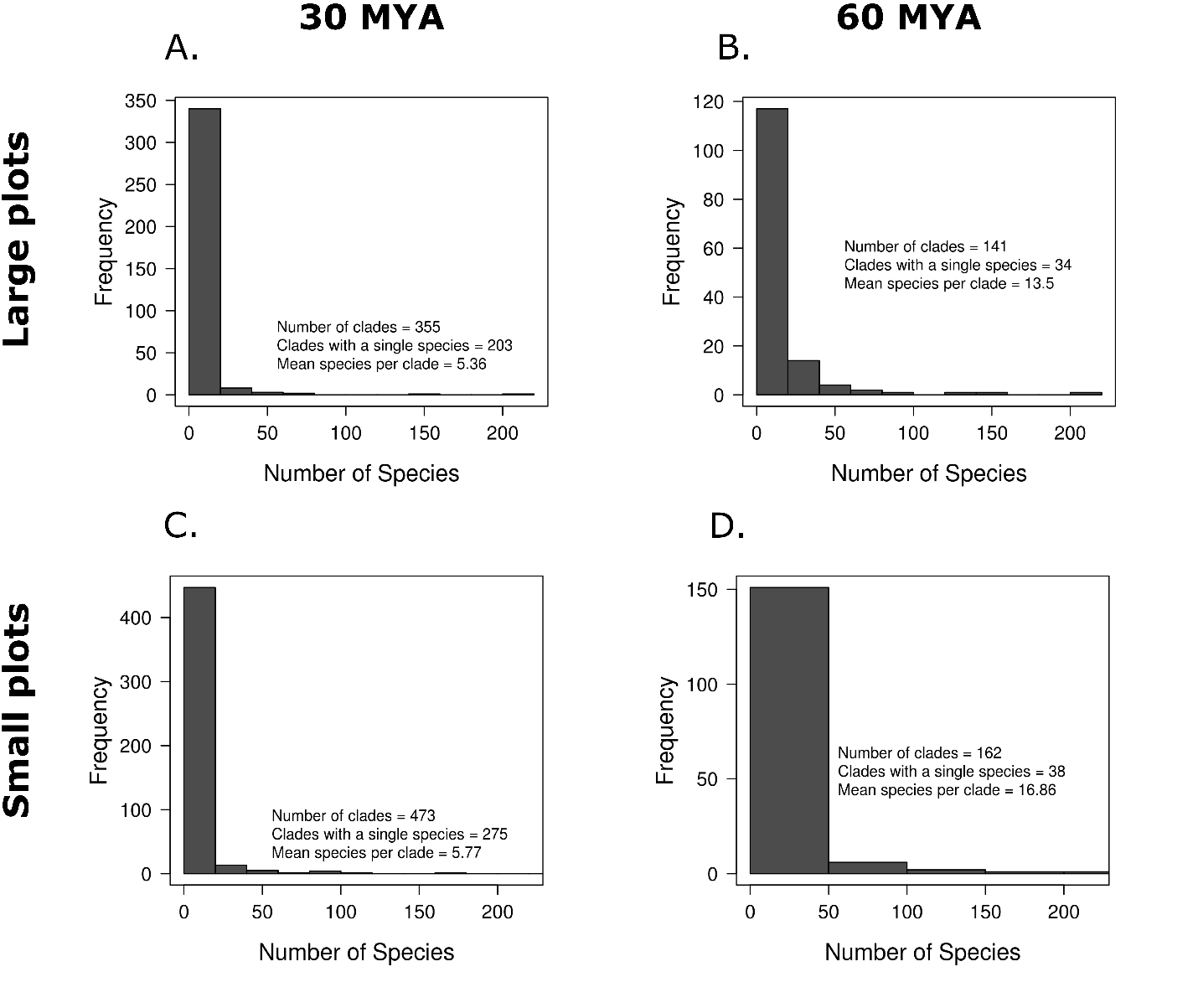

**Figure S2.** **Number of species per clade across datasets.** Clades are ranked in the x-axis from most to least number of species. Panels depict clades present in large plots **(A. & B.)** and small plots **(C. & D.)** for 30 and 60 MY clades. Red dashed line indicates point at which number of species per clade becomes one.

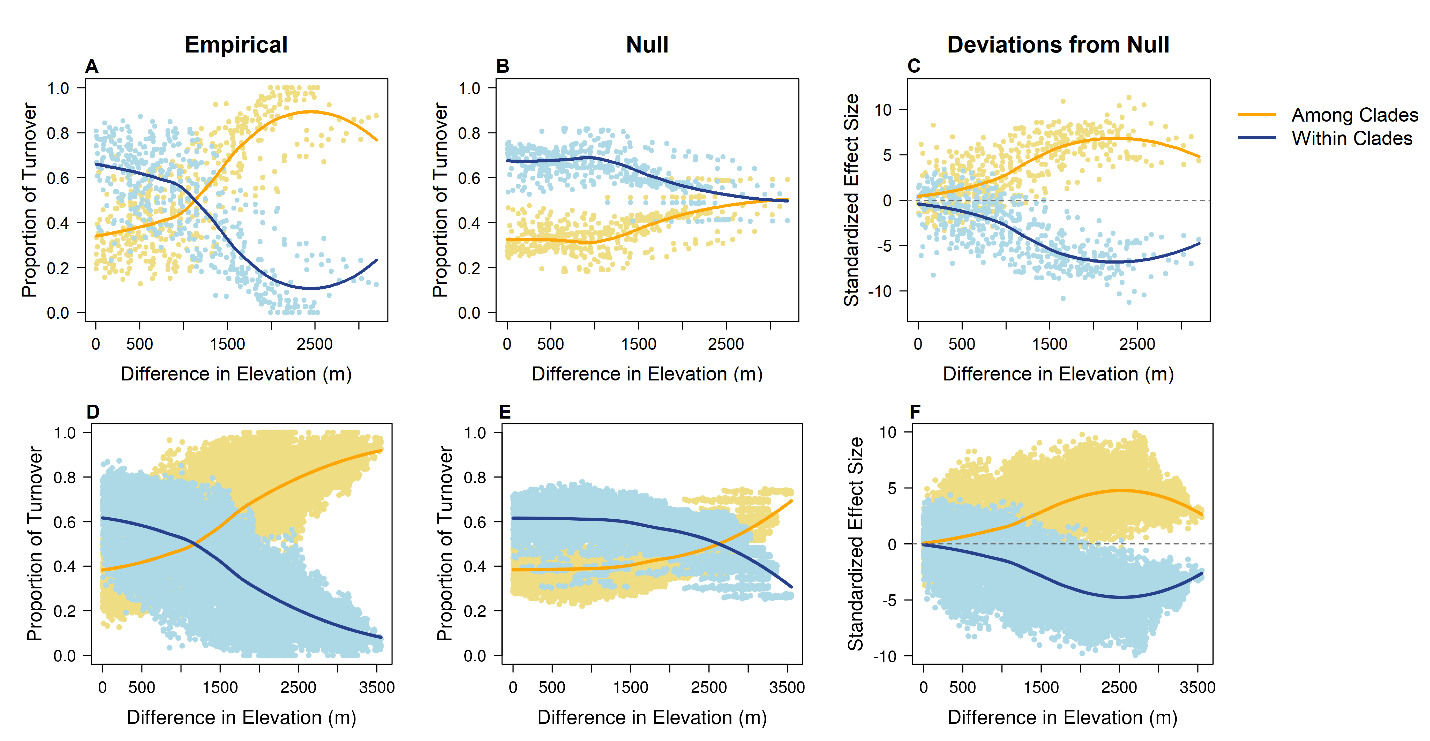

**Figure S3. Decomposition of species turnover across elevational distance into among-clade and within-clade components – 60 MY clades from small and large plots including morphospecies.** Sørensen dissimilarities between each pair of plots were decomposed into within-clades (blue lines) and among-clades (yellow lines) proportions. We then plotted these components of turnover against elevational distance in large plots (first row) and small plots (second row). Finally, we compared **(A&D)** observed patterns in variation of these components with a **(B&E)** tip-randomization null model that removes any phylogenetic structure in the distribution of species across elevational distance, resulting in **(C&F)** standardized effect sizes showing the deviation of the empirical values from null expectations.

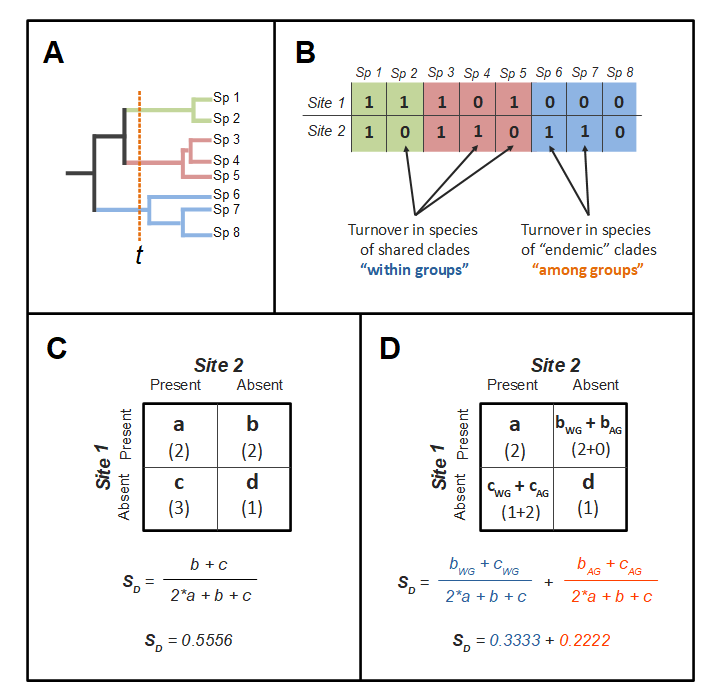

**Figure S4. Decomposition of Sørensen dissimilarity between two sites into within- and among-group components.** **(A)** A putative phylogeny that groups eight species into three clades that diverged from one another before time t. These clades provide a grouping scheme for species, but any other criteria to group species could be used. **(B)** A putative community composition table showing the presences (1s) and absences (0s) of species across two different sites. Cells are colored by species according to the clades (groups) that they belong to. Species contribute to turnover when they are present in a site but missing from the other. Of the three clades, the first two (green and red) have representatives in both sites. The last clade (blue) is endemic to the second site only. Thus, species in the first two clades contribute to within-clade turnover, while species in the last clade contribute to among clade turnover. **(C)** 2x2 frequency table summarizing the number of species present in both sites (a), present in only one site (b and c), or absent from both sites (d). The values corresponding to the table in (B) are presented between parentheses. These numbers are used to calculate the Sørensen dissimilarity index as described by the formula. For our example in (B), the result is 0.556. **(D)** 2x2 frequency table where the b and c values have been partitioned into species within shared clades (b_WG_ and c_WG_) or species among unique clades (b_AG_ and c_AG_). In this way, the Sørensen dissimilarity can be partitioned in to two additive components that correspond to turnover within- and among-clades. For our example in (B), the within-clade component is 0.333 and the among-clade component is 0.222.

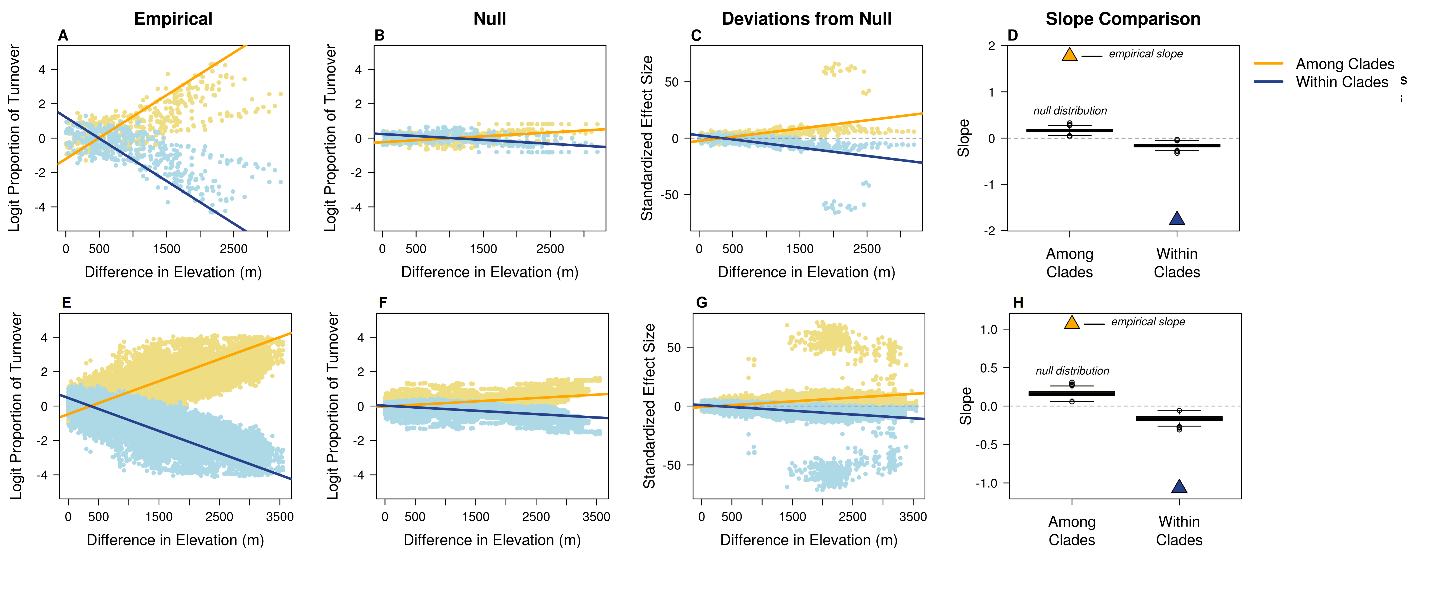

**Figure S5.** **Logit transformed decomposition of species turnover across elevational distance into among-clade and within-clade components – 30 MY clades from small and large plots, including morphospecies.** Sørensen dissimilarities between each pair of plots were decomposed into within-clades (blue lines) and among-clades (yellow lines) proportions and logit transformed. We then plotted these components of turnover against elevational distance in large plots (first row) and small plots (second row). Finally, we compared spatial patterns in variation of these components with a tip-randomization null model that removes any phylogenetic structure in the distribution of species across geography. **(A & E)** empirical patterns; **(B & F)** patterns for the mean of the expectations in the null model; **(C & G)** patterns for standardized effect sizes showing the deviation of the empirical values from null expectations; **(D & H)** comparison of empirical linear slopes for the empirical patterns (stars) with the distribution of 1,000 slopes produced by the null model (boxplots).

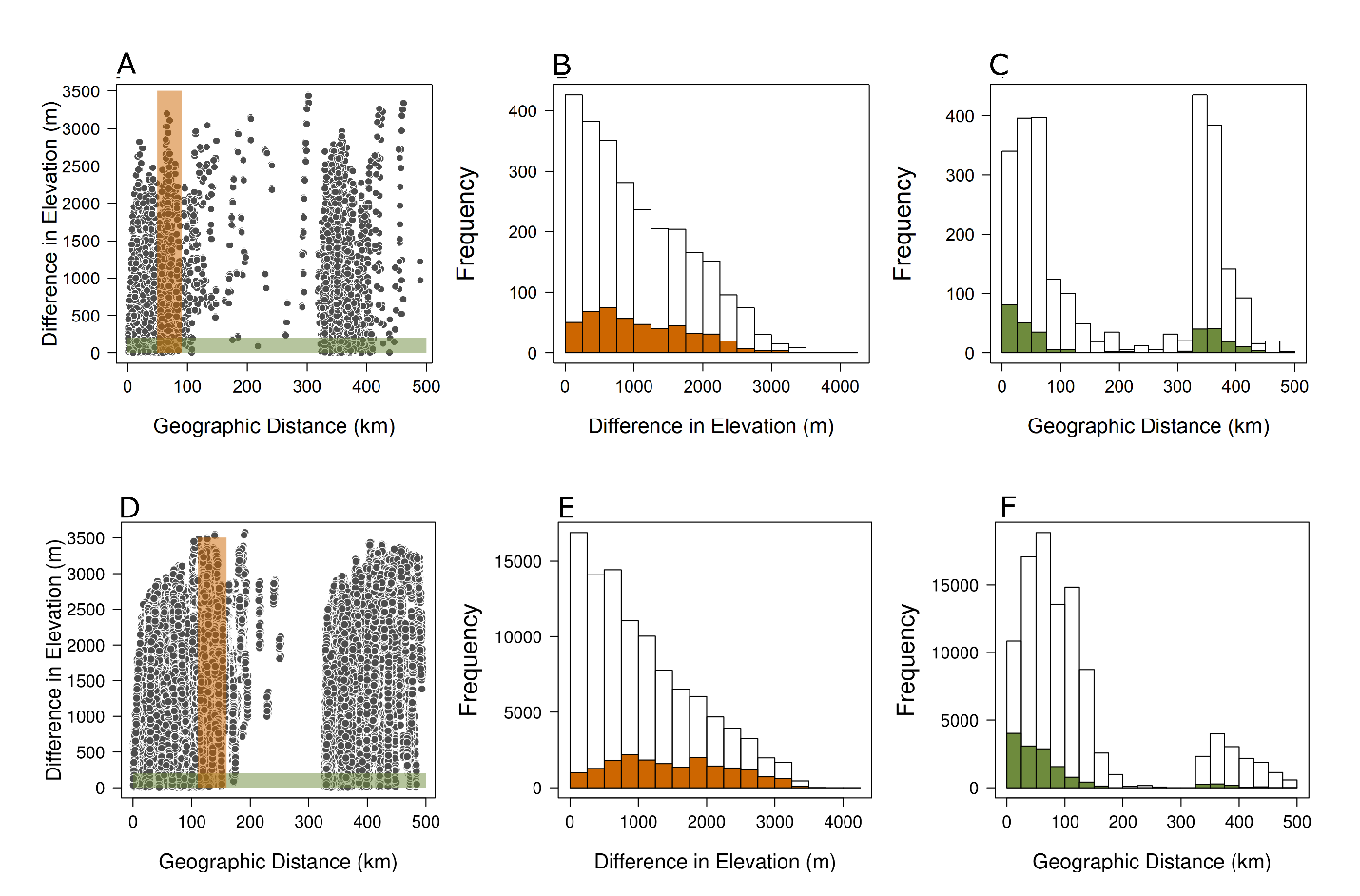

**Figure S6.** **Relationship between difference in elevation and geographic distance in our datasets.** Graphs depict pairwise plot comparisons selected for analyses out of the total number observation in large plots **(first row; A-C)** and small plots **(second row; D-F)**. The objective of this selection was to minimize the effect of geographic distance when calculating turnover across elevations (**B & E**) and to minimize the effect of elevational distance when calculating turnover across geographic distance (**C & F**). All pairwise observations are represented by the grey symbols and the white bars in the histograms. The subset of observations used to study elevation effects are highlighted in orange; while the subset of observations to study geographic distance effects are in green.

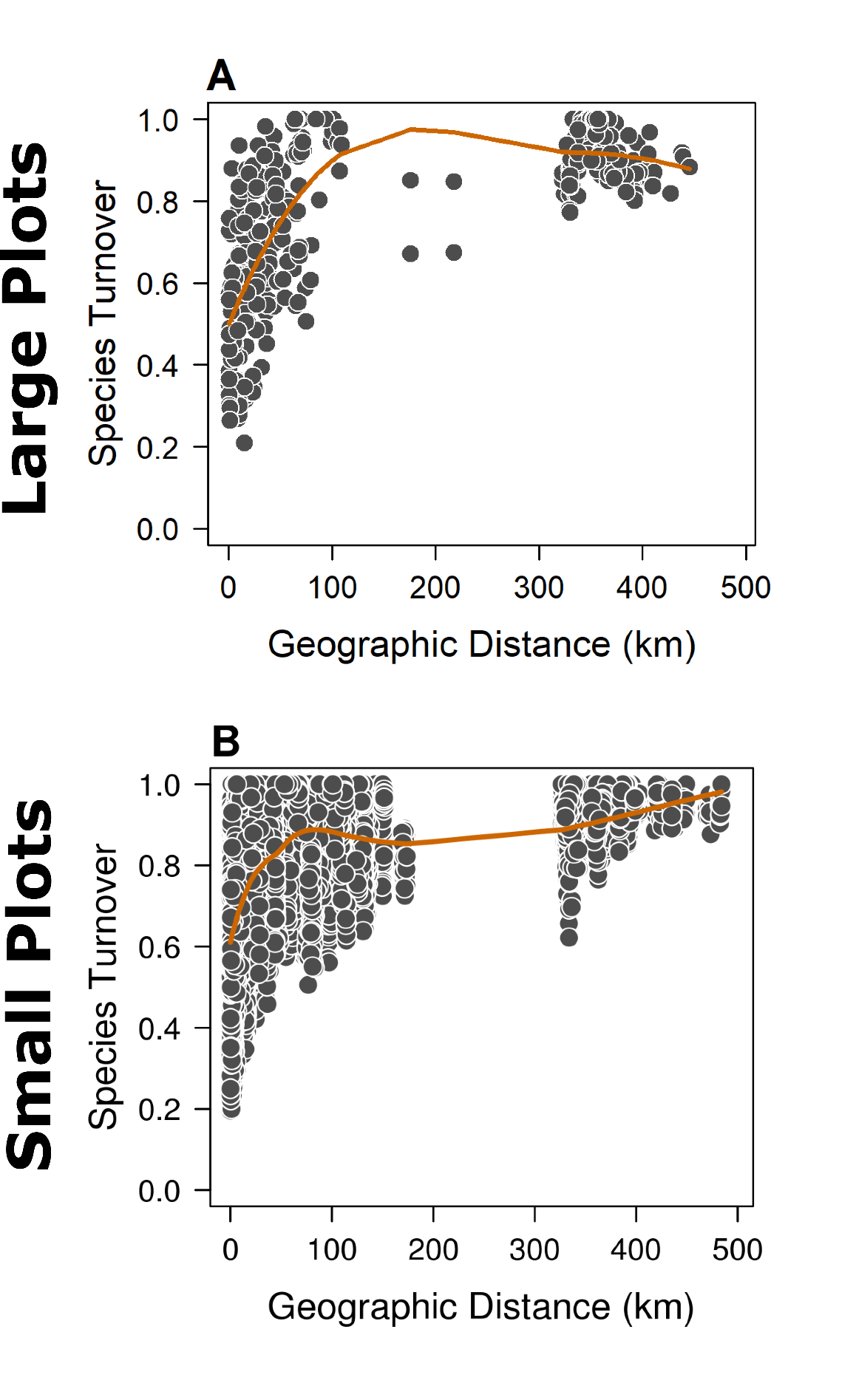

**Figure S7.** **Species turnover across geographic distance.** Sørensen dissimilarity indices plotted against geographic distance for each pair of plots in our two datasets. These patterns are presented separately for **(A)** large 1-ha plots and **(B)** small 0.1-ha plots.

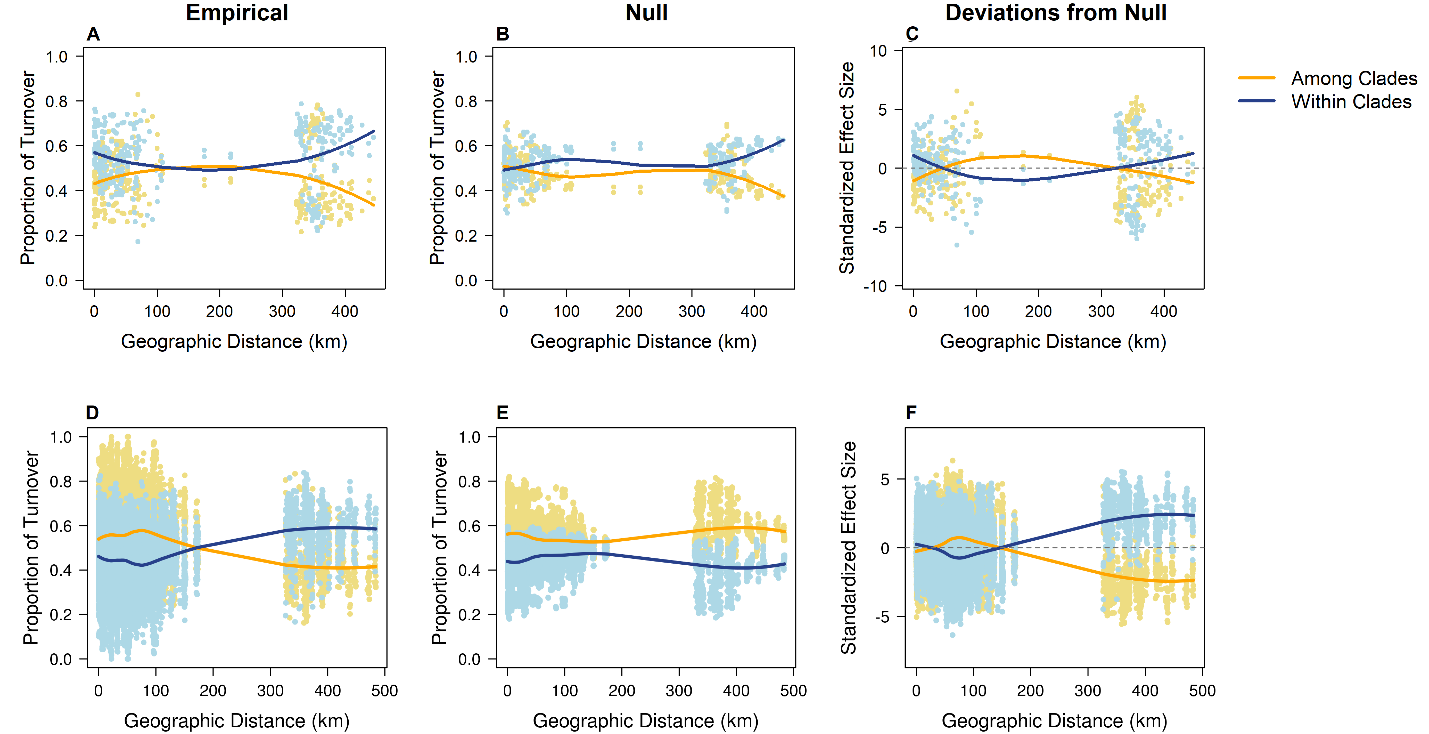

**Figure S8. Decomposition of species turnover across geographic distance into among-clade and within-clade components – 30 MY clades from small and large plots including morphospecies.** Sørensen dissimilarities between each pair of plots were decomposed into within-clades (blue lines) and among-clades (yellow lines) proportions. We then plotted these components of turnover against geographic distance in large plots (first row) and small plots (second row). Finally, we compared **(A&D)** observed patterns in variation of these components with a **(B&E)** tip-randomization null model that removes any phylogenetic structure in the distribution of species across elevational distance, resulting in **(C&F)** standardized effect sizes showing the deviation of the empirical values from null expectations.
